# Population and individual effects of non-coding variants inform genetic risk factors

**DOI:** 10.1101/065144

**Authors:** Pala M., Z. Zappala, M. Marongiu, X. Li, J.R. Davis, R. Cusano, F. Crobu, K.R. Kukurba, F. Reiner, R. Berutti, M.G. Piras, A. Mulas, M. Zoledziewska, M. Marongiu, F. Busonero, A. Maschio, M. Steri, C. Sidore, S. Sanna, E. Fiorillo, A. Battle, J. Novembre, C. Jones, A. Angius, G.R. Abecasis, D. Schlessinger, F. Cucca, S.B. Montgomery

**Affiliations:** Istituto di Ricerca Genetica e Biomedica (IRGB), CNR, Monserrato, Italy; Department of Pathology, Stanford University School of Medicine; Department of Genetics, Stanford University School of Medicine; CRS4, Advanced Genomic Computing Technology, Pula, Italy; Center for Computational Biology, John Hopkins University; Department of Human Genetics, University of Chicago; Center for Statistical Genetics, University of Michigan, Ann Arbor, MI; Laboratory of Genetics, NIA, Baltimore, Maryland; Dipartimento di Scienze Biomediche, Università di Sassari, Sassari, Italy

**Author notes:** co-first authors (Pala M., Z. Zappala). co-senior authors (F. Cucca, S.B. Montgomery).

## Abstract

Identifying functional non-coding variants can enhance genome interpretation and inform novel genetic risk factors. We used whole genomes and peripheral white blood cell transcriptomes from 624 Sardinian individuals to identify non-coding variants that contribute to population, family, and individual differences in transcript abundance. We identified 21,183 independent expression quantitative trait loci (eQTLs) and 6,768 independent splicing quantitative trait loci (sQTLs) influencing 73 and 41% of all tested genes. When we compared Sardinian eQTLs to those previously identified in Europe, we identified differentiated eQTLs at genes involved in malarial resistance and multiple sclerosis, reflecting the long-term epidemiological history of the island’s population. Taking advantage of pedigree data for the population sample, we identify segregating patterns of outlier gene expression and allelic imbalance in 61 Sardinian trios. We identified 809 expression outliers (median z-score of 2.97) averaging 13.3 genes with outlier expression per individual. We then connected these outlier expression events to rare non-coding variants. Our results provide new insight into the effects of non-coding variants and their relationship to population history, traits and individual genetic risk.

## INTRODUCTION

Human migration and rapid population expansion have led to an abundance of population and individual-specific genetic variants (Coventry et al., 2010; Nelson et al., 2012; Tennessen et al., 2012; 1000 Genomes Project Consortium et al., 2015; UK10K Consortium et al., 2015). Within protein-coding regions of the genome, multiple studies have identified numerous rare alleles and have reported on the impact of loss-of-function alleles (MacArthur et al., 2012; Flannick et al., 2014; Exome Aggregation Consortium et al., 2015; Li et al., 2015; Sulem et al., 2015; Narasimhan et al., 2016), particularly in isolated populations (Moltke et al., 2014). However, protein-coding variation represents only a fraction of the genetic variation that can impact human traits. Large studies of gene expression have greatly advanced our ability to identify functional variation in the non-coding portions of the genome (Battle et al., 2014; GTEx Consortium, 2015; Lappalainen et al., 2013), and many of these variants have been connected to common genetic diseases (Maurano et al., 2012; Nicolae et al., 2010). However, few studies to date have had access to whole genome sequencing data, family relationships and auxiliary complex trait data from research participants. Such data has the potential to empower the assessment of population and individual-specific consequences of non-coding variants.

To overcome this, we sequenced RNA from white blood cells of 624 individuals from the founder population of Sardinia. The Sardinian population has several advantages: their DNA includes the bulk of mainland European DNA variation, but due to a period of relative isolation for >10,000 years, many alleles have been added, and many old and novel variants have reached dramatically higher frequencies which should improve power to detect associations between those variants and traits such as gene expression (Lim et al., 2014; Moltke et al., 2014; Peltonen et al., 2000; Sidore et al., 2015). In addition, the SardiNIA study cohort has been extensively genotyped and phenotyped and consist of both unrelated and related individuals (Orrù et al., 2013). By combining RNA-seq data with whole genome sequence data, we discover expression and splicing quantitative trait loci (e/sQTLs) that are specific to the isolated Sardinian population. As this is the first study to integrate both whole genomes and transcriptomes from multiple families, we developed a framework that leverages these family relationships in order to identify the regulatory impact of rare non-coding variants. We identify extreme gene expression outliers that segregate within these families and investigate the distribution and associated functional annotations of putatively causal rare variants as well as their influence on individual disease risk

## RESULTS

### Expression and splicing quantitative trait discovery in Sardinia

The 624 participants, all from four towns in the Lanusei Valley in the Ogliastra region of Sardinia, were enrolled from a cohort of 6,921 in the SardiNIA longitudinal study of aging (Pilia et al., 2006) (see Methods). The entire SardiNIA cohort was genotyped using two custom Illumina arrays, the Cardio-MetaboChip and ImmunoChip. A subset of 1,146 Sardinians were additionally whole-genome sequenced at low coverage (average four-fold) as previously described (Orrù et al., 2013; Sidore et al., 2015), producing an integrated map of ∼15 million SNPs. For RNA, we sequenced a median of ∼59 million 51-bp paired-end reads per participant (over 36 billion reads in total). After quantification and quality control 15,243 and 12,603 genes were sufficiently expressed for eQTL and sQTL analyses, respectively (see Methods; Table 1). To account for confounders that can reduce power to discover *cis-*QTLs, we applied hidden factor correction using PEER (Stegle et al., 2012). We were able to identify and remove confounders due to gender, age, blood cell counts, and sequencing (see Methods).

**Table 1.** Expression traits with at least one eQTL. We report the number of tests performed and the number of significant QTL associations for different expression traits at a false discovery rate of 5%. Associations that are significant by BH are significant after Bonferroni correction and Benjamini-Hochberg adjustment (see Methods)

| Measurement | Trait type | # of traits tested | # of traits with at least one QTL (FDR 5%) |  |
| --- | --- | --- | --- | --- |
|  |  |  | BH | By permutation |
| Gene-level | Protein coding | 11477 | 8381 (73%) | 9019 (79%) |
|  | lncRNA | 1694 | 991 (69%) | 1258 (74%) |
|  | miRNA precursors | 172 | 48 (27%) | 55 (32%) |
|  | Other | 1900 | 935 (39%) | 835 (44%) |
|  | Total | 15243 | 10329 (68%) | 11167 (73%) |
| Isoform-proportion* | Protein coding | 11116 | 3865 (35%) | 4515 (41%) |
|  | lncRNA | 826 | 335 (41%) | 373 (45%) |
|  | Other | 661 | 213 (32%) | 323 (49%) |
|  | Total | 12603 | 4413 (35%) | 5120 (41%) |

(\*) Isoforms results are reported at gene-level (only one sQTL per gene is reported)

To discover eQTLs, we tested the association of genotype with expression level for all variants within ±1MB of a target gene’s transcription start site (TSS) for all individuals with genetic data from the integrated map (N = 606). At a false discovery rate (FDR) of 5%, we identified eQTLs for the majority of tested genes (Table 1). We next applied conditional analyses to characterize the number of independent eQTLs per gene (see Methods). We found that approximately half of all protein-coding and lncRNA transcripts were influenced by at least two independent eQTLs; miRNAs, however, were mostly associated with a single eQTL (Table 2). At the extreme, we found that a single protein-coding gene, *ITGB1BP1,* was affected by 14 independent eQTLs. *ITGB1BP1* encodes an integrin binding protein that is implicated in upstream regulation of immune-critical TNF/NF-kB transcriptional regulation. We also identified *NBPF1*, a lncRNA of unknown function, that was affected by 11 independent eQTLs (Table S5). In total, we mapped at least one eQTL for 73% of tested genes, corresponding to 11,167 different eQTLs. After conditional analyses, we identified an extra 10,016 eQTLs for a total of 21,184 eQTLs (Table 2).

**Table 2.** Independent QTLs segmented by gene type. We report the number of independent QTLs for gene-level and isoform-level analyses. Isoform results are grouped by their respective gene.

| Max # of independent QTLs | Gene-level (# of genes) |  |  | Isoform-proportion (# of genes) |  |
| --- | --- | --- | --- | --- | --- |
|  | Protein coding | lncRNA | miRNA precursors | Protein coding | lncRNA |
| 1 | 4215 | 598 | 44 | 3489 | 281 |
| 2 | 2833 | 386 | 8 | 799 | 60 |
| 3 | 1235 | 170 | 2 | 165 | 22 |
| 4 | 428 | 66 | 0 | 36 | 8 |
| 5 | 175 | 18 | 1 | 18 | 1 |
| 6 | 82 | 6 | 0 | 3 | 1 |
| 7 | 27 | 5 | 0 | 5 | 0 |
| 8 | 14 | 3 | 0 | 0 | 0 |
| ≥9 | 10 | 6 | 0 | 0 | 0 |
| Total | 9019 | 1258 | 55 | 4515 | 373 |

To discover sQTLs, we tested the association of genotype with the ratio of known transcript abundances calculated using Cufflinks (Trapnell et al., 2010). At an FDR of 5%, we observed significant sQTLs for nearly half of tested protein-coding genes and lncRNAs. In total, this is over a thousand more sQTLs than previously reported (Battle et al., 2014; Lappalainen et al., 2013). In comparison to eQTLs, we found that protein-coding genes and lncRNAs were less likely to have multiple independent sQTLs (Table 2); however, we found five protein-coding genes that were influenced by as many as seven independent sQTLs (Table S6). Notably, two of these genes affect transcription and splicing itself, and are expected to impact the immune system. These genes include: *POLR2J2*, one of two nearly identical polymerase II subunit genes known to produce alternative transcript; and *SMN1*, which functions in the assembly of the spliceosome. The other three sQTL genes are directly related to immune function: the nonclassical class I heavy chain paralogue *HLA-G*; the class I heavy chain receptor *HLA-C*; and *ITGB1BP1*. The *ITGB1BP1* gene, which has 8 exons that extend over 16KB and are spliced into 21 isoforms, had extreme numbers of independent eQTLs and sQTLs, suggesting that it is a large mutational target for modulators of expression. While less pervasive than eQTLs, we mapped at least one sQTL for 41% of tested genes, corresponding to 5120 different sQTLs. After conditional analyses, we identified an extra 1,648 sQTLs for a total of 6,768 sQTLs (Table 2). All QTL results are available from http://montgomerylab.stanford.edu/resources/sardinia.html.

### Comparison of Sardinia and European eQTLs identifies novel functional and trait-associated variants

We next measured the replication of Sardinian QTLs with European QTLs in the Depression Genes and Networks (DGN) and GEUVADIS studies (Battle et al., 2014; Lappalainen et al., 2013). For Sardinian eQTLs that were testable in each study, the replication rate was 92% in DGN and 72% in GEUVADIS reflecting the high–degree of sharing of common European alleles within Sardinia (Table 2). For sQTLs, the replication rate was 72% in DGN and 76% in GEUVADIS. Additionally, we tested eQTLs and sQTLs identified in either DGN or GEUVADIS for replication in the Sardinia cohort and found that between 89-92% of eQTLs and 70-97% of sQTLs replicated (Table S4). For 2,568 eQTLs and 1,152 sQTLs in Sardinia, replication could not be tested because the SNPs were absent in Europe or only present at a minor allele frequency below 1%. Of these QTLs, 473 eQTL and 182 sQTL variants were novel in Sardinia when compared to the 1000 Genomes, dbSNP, UK10K and the ExAC database, representing new and/or previously uncaptured functional variation.

To determine if these novel eQTLs were associated with traits measured in Sardinia, we tested all 473 novel eQTL variants for associations with 15 blood cell measurements in the whole SardiNIA cohort (N ≈ 6,000). We identified 5 associations (5 traits and 5 variants, in total) that were significant after correcting for multiple testing (p-value < 7.63e-06). For each association, we then retested the trait association for all variants within ±1MB of the target gene to identify the subset of loci where both the Sardinia-specific eQTL variant and the top trait-associated variant within this region were in high LD (r^2^ > 0.8). We identified a Sardinia-specific eQTL for *ARHGDIB* that was also linked to the top trait associated variant for neutrophil percentage, which is also Sardinia-specific (Supplemental Figure S4; top neutrophil percentage variant chr12:14190223, p-value = 3.80e-06; top eQTL, chr12:15553026, p-value =7.69e-06, r^2^ = 0.86). Within this locus only 3 of 14 variants that passed our LD filter were previously reported outside of Sardinia (allele frequencies in Europe below 0.002). Of note, one of these variants (chr12:15095546, p-value = 3.85e-06, r^2^ with top neutrophil signal = 0.84) is a nonsense allele that had been observed only once in the ExAC database but has a frequency >1% in Sardinia, with the direction of effect on expression consistent with nonsense-mediated decay. The *ARHGDIB* gene presents a biologically plausible target for this association as it is a multi-functional protein with a central role in inhibition of cell migration, and ARHGDIB^-/-^ mice show changes in lymphocyte expansion and survival in culture (Dovas and Couchman, 2005).

### Sardinia eQTLs exhibit founder population effects and evidence of selection

As genetic analyses in founder populations like Sardinia are expected to have increased statistical power based on relatively low heterogeneity and shared environment (Peltonen et al., 2000), we compared the observed impact of Sardinian eQTLs to European eQTLs. Using an identical pipeline and controlling for various differences in study parameters, we regenerated European eQTLs from the DGN and GEUVADIS studies (see Methods). When comparing eQTLs between these studies and Sardinia, we observed increased correlation between expression and genotype for Sardinian eQTLs (Figure 1A). This could reflect founder population effects or reduced technical noise in our study. As allele-specific expression signals have been demonstrated to be more robust to technical noise levels, we also compared Sardinian allele-specific expression QTLs (aseQTLs) to European aseQTLs (Castel et al., 2015). Consistent with a founder population effect, we observed an increased correlation of genotype and allelic expression for Sardinia aseQTLs (Figure 1B).

**Figure 1.**
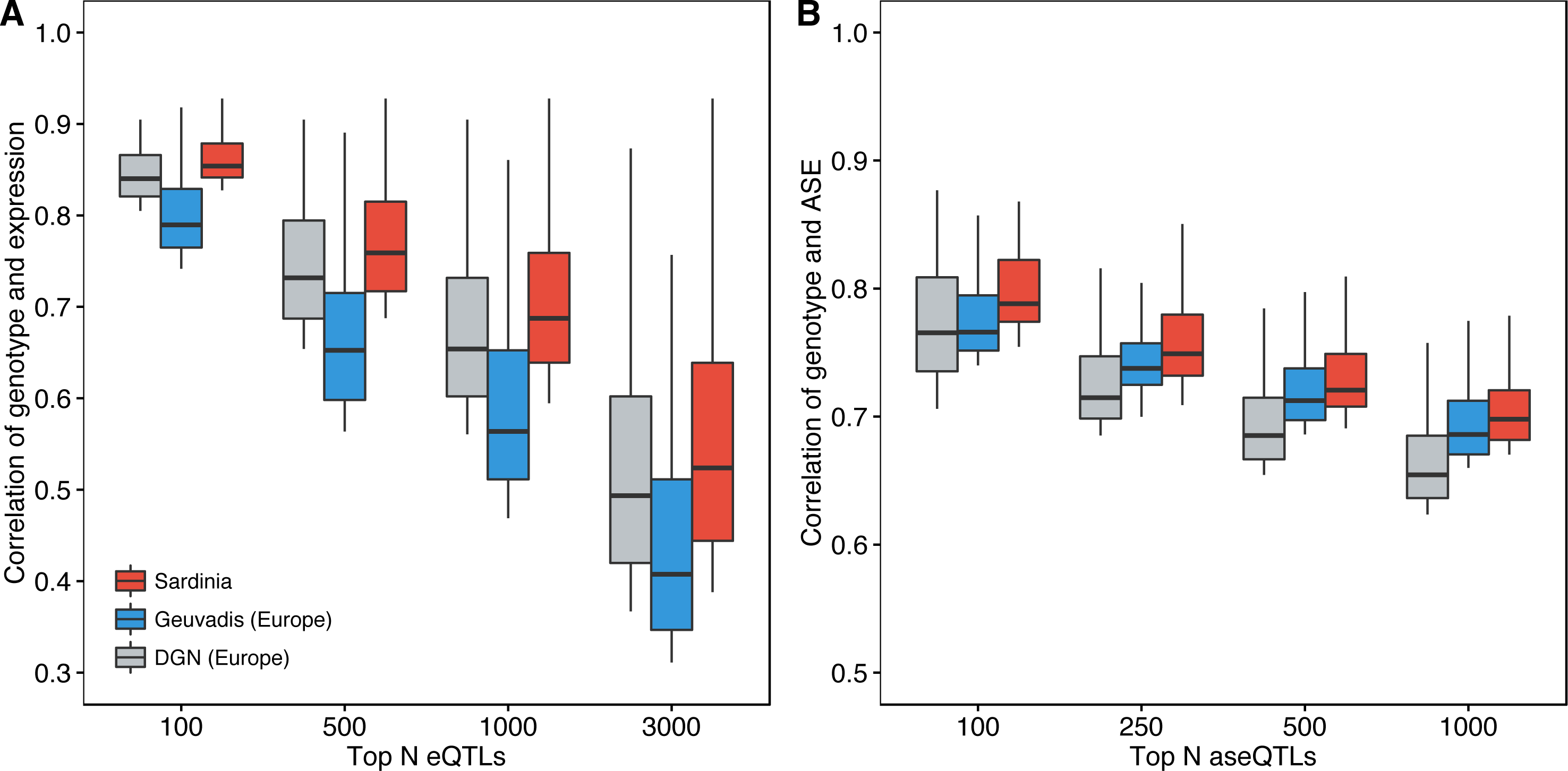
QTLs show large effect sizes in Sardinia compared to Europe. **(a)** eQTLs are compared for the top eQTLs found in Sardinia, Geuvadis and DGN studies demonstrating increased correlation between genotype and expression level for Sardinian eQTLs. **(b)** Allele-specific expression QTL (aseQTLs) are compared for the top aseQTLs in Sardinia, Geuvadis and DGN studies demonstrating increased correlation between genotype and expression level for Sardinian aseQTLs. To make analyses comparable for both eQTLs and aseQTLs, 188 unrelated individuals from each study were uniformly processed and the intersection of quantifiable genes was selected for QTL discovery.

To identify eQTLs where founder effects, genetic drift, or selective pressure have significantly influenced the prevalence of these alleles in Sardinia, we compared the Sardinian allele frequencies of eQTLs and sQTLs with the corresponding European allele frequency reported by the 1000 Genomes Project (1000 Genomes Project Consortium et al., 2015). We found that 11% of significant eQTLs were differentiated at an allele frequency greater than 10% (Figure 2A). In addition, we observed longer tracts of linkage disequilibrium (LD) decay in Sardinians conditioned on the extent of allelic differentiation (Figure 2B). Ten of these LD tracts showed evidence of hard selective sweeps (integrated haplotype scores (iHS) > 2.5), consistent with a proportion of these eQTLs having undergone recent positive selection (Voight et al., 2006).

**Figure 2.**
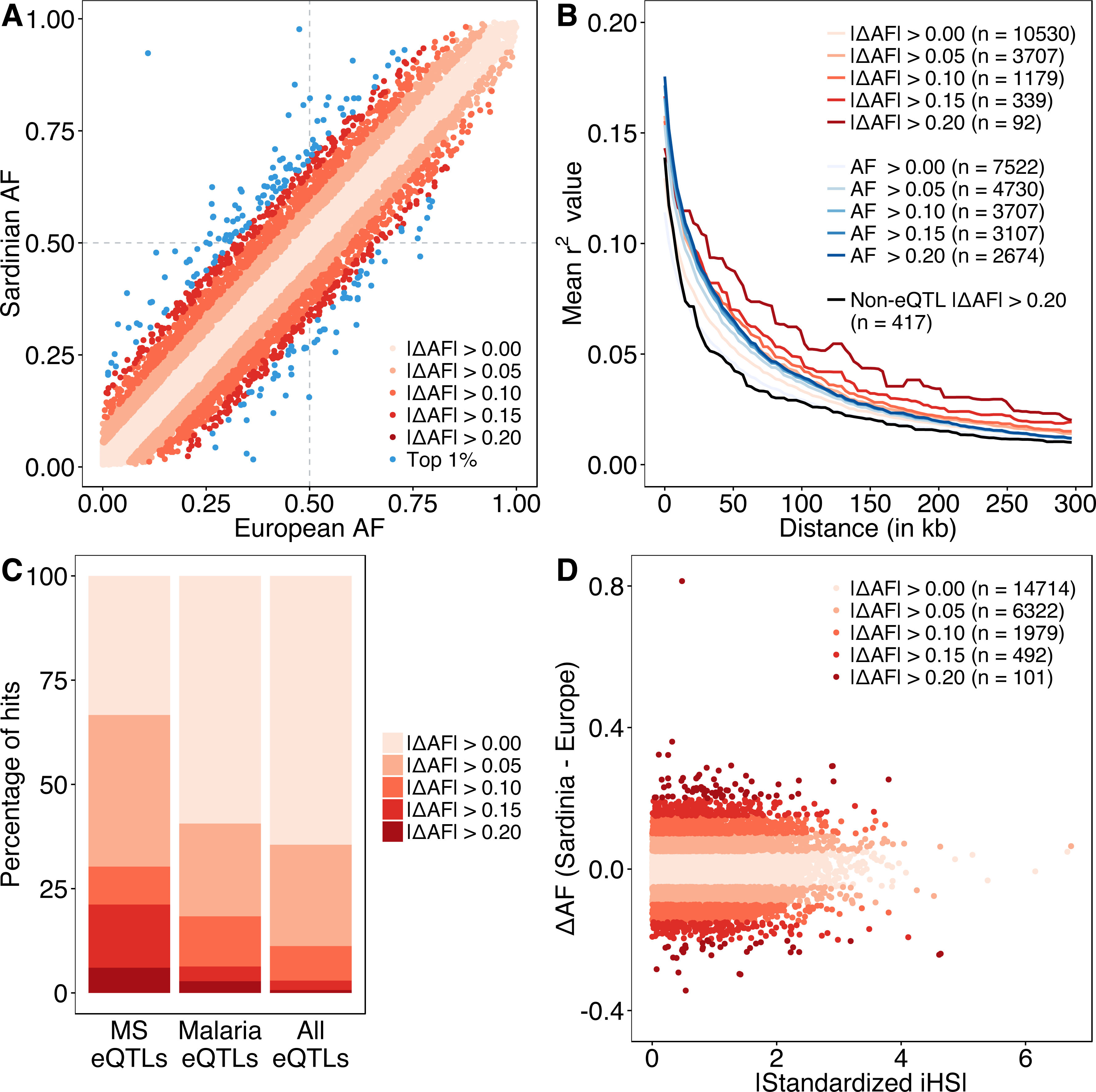
Differentiated eQTLs in Sardinia. **(a)** Sardinian eQTLs are plotted compared to their allele frequency differences between Europe (measured from the 1000 Genomes Project) and Sardinia. Sample sizes: |ΔAF| > 0.00 (N = 19108 eQTLs), > 0.05 (N = 6793), > 0.10 (N = 2151), > 0.15 (N = 567), > 0.20 (N = 134), and Top 1% (N = 192) **(b)** eQTLs with larger allele frequency differences compared to Europe have larger tracts of LD decay as evidence for recent positive selection. These are compared to eQTLs which have comparable allele frequencies in Sardinia and minimal (+/- 2.5%) changes in allele frequencies compared to Europe (blue lines). They are also compared to randomly selected non-eQTL variants with large allele-frequency changes (black line). **(c)** eQTLs linked to multiple sclerosis variants and malaria genes are both enriched in allele frequency difference changes between Sardinia and Europe. **(d)** Multiple eQTLs have large differences in iHS and allele frequency informing recent positive selection.

### Highly differentiated eQTLs are enriched for malaria and multiple sclerosis genes

We next tested whether two epidemiological factors present in Sardinia were reflected among highly differentiated eQTLs. Until the mid-twentieth century, the Sardinian population suffered high mortality rates due to malaria (Kaneko et al., 2000; Tognotti, 2009), and continues to have a higher prevalence of multiple sclerosis (MS) relative to other Caucasian populations in the Mediterranean basin (Pugliatti et al., 2006; 2002). Indeed, we identified a significant enrichment for known malarial resistance genes (p-value = 0.0015) and genes associated with MS (EBI/NHGRI GWAS Catalog p-value = 1.84 x 10^−5^ and ImmunoBase p-value = 1.17 x 10^−8^) among the top 1% of differentiated eQTLs (mean allele frequency difference of ∼17%) (Figure 2C, Table S3). MS had the highest enrichment among 354 traits tested from the EBI/NHGRI GWAS catalog and among 19 traits tested from the ImmunoBase catalog. Furthermore, one of the most differentiated eQTLs was the *BAFF* gene (p-value = 8.051e-12, ΔAF*SRD-EUR* = 0.25), which is known to be involved in response and survival to malaria infection (Liu et al., 2012; Scholzen and Sauerwein, 2013; Scholzen et al., 2014) and has unique evolutionary history in Sardinia (Steri *et al,*submitted) (Figure 2D). We also observed relevant variants in the *CR1* gene which is involved in complement activation and immune complex formation during malaria infection (Kosoy et al., 2011; Stoute, 2011). *CR1* has two eQTLs (chr1:207275799 and chr1:207667190) and 9 sQTLs. The eQTL at chr1:207667190 is highly differentiated between Sardinia and Europe (ΔAF*SRD-EUR* = −0.25).

Among the 9 sQTLs associated with *CR1*, two of them are highly differentiated: chr1:207681501 (ΔAF*SRD-EUR* = 0.42) influences abundance of ENST00000367051 and chr1:207716099 (ΔAF*SRD-EUR* = 0.43) influences abundance of ENST00000529814. Both sQTLs are tightly linked and in high LD (r^2^ = 0.99 and 0.95) with a chr1:207757515 variant previously identified to be associated with erythrocyte sedimentation rate in the SardiNIA cohort (Naitza et al., 2012).

### Heritable patterns of extreme gene expression in families

Beyond the unique history of the Sardinia population, the availability of family relationship data in the SardiNIA sample provided an opportunity to identify the impact of rare, large-effect non-coding variation. Specifically, we developed a likelihood ratio test to identify patterns of extreme gene expression that segregated in families (Figure 3A; see Methods). We tested 61 Sardinian trios for the 15,243 genes included in our eQTL analyses and identified 809 genes where a parent and child are both expression outliers (median z-score = 2.97) at an FDR of 10% (Figure 3B). On average we found 13.3 shared gene expression outliers per trio.

**Figure 3.**
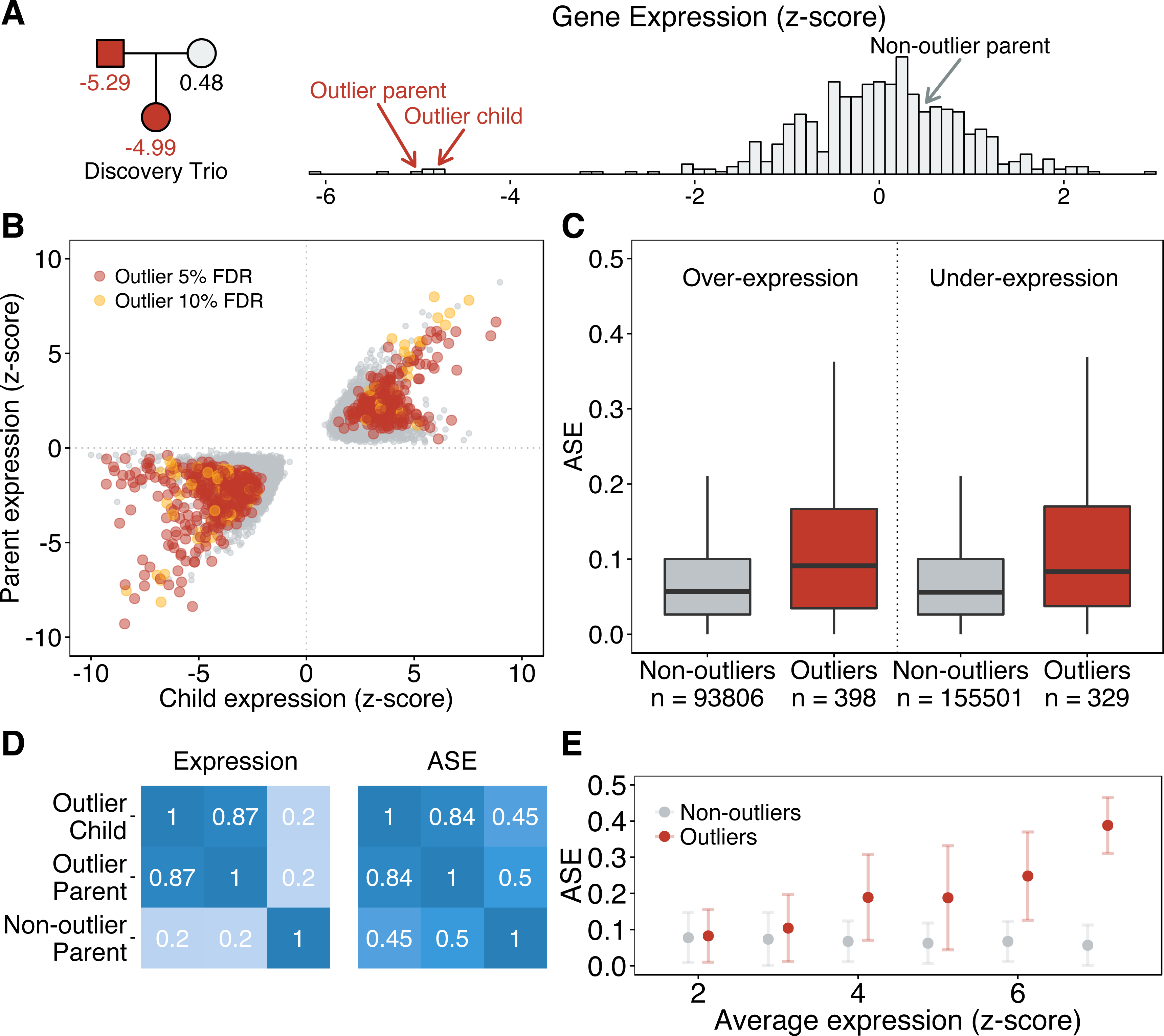
Outlier gene expression in Sardinian trios. **(a)** An example of a significant gene expression outlier effect that segregates in a single Sardinia trio. The father and daughter both under express the *RINL* gene and share a rare splicing variant. **(b)** A scatterplot showing the sharing of extreme gene expression patterns between parents and children in 61 Sardinian trios, with significant outliers highlighted (orange = 5% FDR and yellow = 10% FDR). **(c)** Heterozygous sites in outlier genes show elevated levels of allele-specific expression in outlier individuals (red) versus the rest of the population (gray). **(d)** Correlation matrices for gene expression and allele-specific expression within outlier trios suggest that the extreme regulatory effects are restricted to the affected individuals and not primarily a family-specific event due to a shared environment. **(e)** The relationship between outlier gene expression and the degree of allele-specific expression observed in outliers (red) and non-outliers (gray).

Several lines of evidence suggest shared expression outliers are not due simply to parent-offspring shared environment. There was little correlation of gene expression between the outlier parent and the non-outlier partner (Pearson r = 0.20) (Figure 3D). Additionally, mothers and fathers were equally likely to be the outlier parent (p = 0.20), regardless of the sex of the child (p = 0.83) (Supplemental Figure S8). We found almost twice as many shared under-expression outliers (529 outliers, 65%) as over-expression outliers (280 outliers, 35%), consistent with observations of the effects of random substitutions in promoters and enhancers in massively parallel reporter assays (Kwasnieski et al., 2012; Melnikov et al., 2012; Patwardhan et al., 2012). Further, since rare variants tend to be heterozygotic and thus only influence on allele, we hypothesized that outlier parents and children would be enriched for allele-specific expression compared to non-outlier controls. We found that allele-specific expression was significantly enriched in outlier individuals for both under- and over-expression outliers (adjusted Wilcoxon rank-sum p-value = 6.0e-6) (Figure 3C). This is likely a conservative estimate of the true enrichment, given the inherently low levels of read depth in under-expression outliers that limits the ability to measure allelic effects in outlier genes. These allelic effects were consistent between outlier parents and children (Pearson r = 0.84) but not with the other, non-outlier parent (Figure 3D). The strength of the outlier effect was also significantly associated with the enrichment of allele-specific expression (Spearman ρ = 0.338, p-value < 1e-6), reflecting the capacity of allele-specific effects to impact total expression (Figure 3E).

### Rare variants can drive extreme gene expression and contribute to individual genetic risk factors

Using the combination of whole genome data and family relationships, we were able to characterize potential causal variants underlying expression outliers. We first identified 3464 rare variants (Sardinia MAF < 1%) that were located in 250KB windows adjacent to the transcription start site (TSS) and end site (TES) of outlier genes and were unambiguously transmitted from the outlier parent to the outlier child. We also identified an equivalent set of 245,165 rare variants in the same genomic loci that were unambiguously transmitted between non-outlier parents and children. We found at least one shared rare variant for 509 of the outlier genes (63%), with an average of 6.8 variants shared by outliers versus 4.0 shared by non-outliers (enrichment = 1.71, 95% confidence interval 1.65 - 1.77). Of interest, rare variants shared by outlier individuals were concentrated within 5KB of the TSS (enrichment = 3.61, 95% confidence interval 2.96 – 4.24) and TES (enrichment = 3.00, 95% confidence interval 2.44 – 3.54) (Figure 4A) of outlier genes, similar to what has been observed for common regulatory variation (Veyrieras et al., 2008). Furthermore, rare variants shared by outliers were enriched in multiple functional annotations (Figure 4B). For variants in the 50KB window adjacent to the TSS, this enrichment was most notable in splice donor/acceptor sites (log odds = 4.05, p-value = 2.52e-07) and regions associated with active transcription, including promoters (log odds = 0.91, p-value = 8.8e-09) and enhancers (log odds = 0.42, p-value = 0.0094) (enrichment data for different genomic window sizes is provided in Supplementary Tables S10-11)(Roadmap Epigenomics Consortium et al., 2015). We further investigated whether other carriers of these variants displayed the same outlier expression profile as the parent-child pairs. We analyzed 2,912 variants (84% of the 3,464 outlier variants) that were heterozygous in at least four individuals in the cohort, regressing outlier gene expression on genotype at the rare variant position. The largest and most significant of these genotype-expression associations for both over- and under-expression outliers were concentrated at the TSS of outlier genes (Figure 4C). Additionally, we found that metrics of conservation (GERP, PhyloP) and predicted functional relevance (FitCons, CADD) all discriminated the most significant associations (Figure 4D) (Cooper et al., 2005; Gulko et al., 2015; Kircher et al., 2014; Pollard et al., 2010).

**Figure 4.**
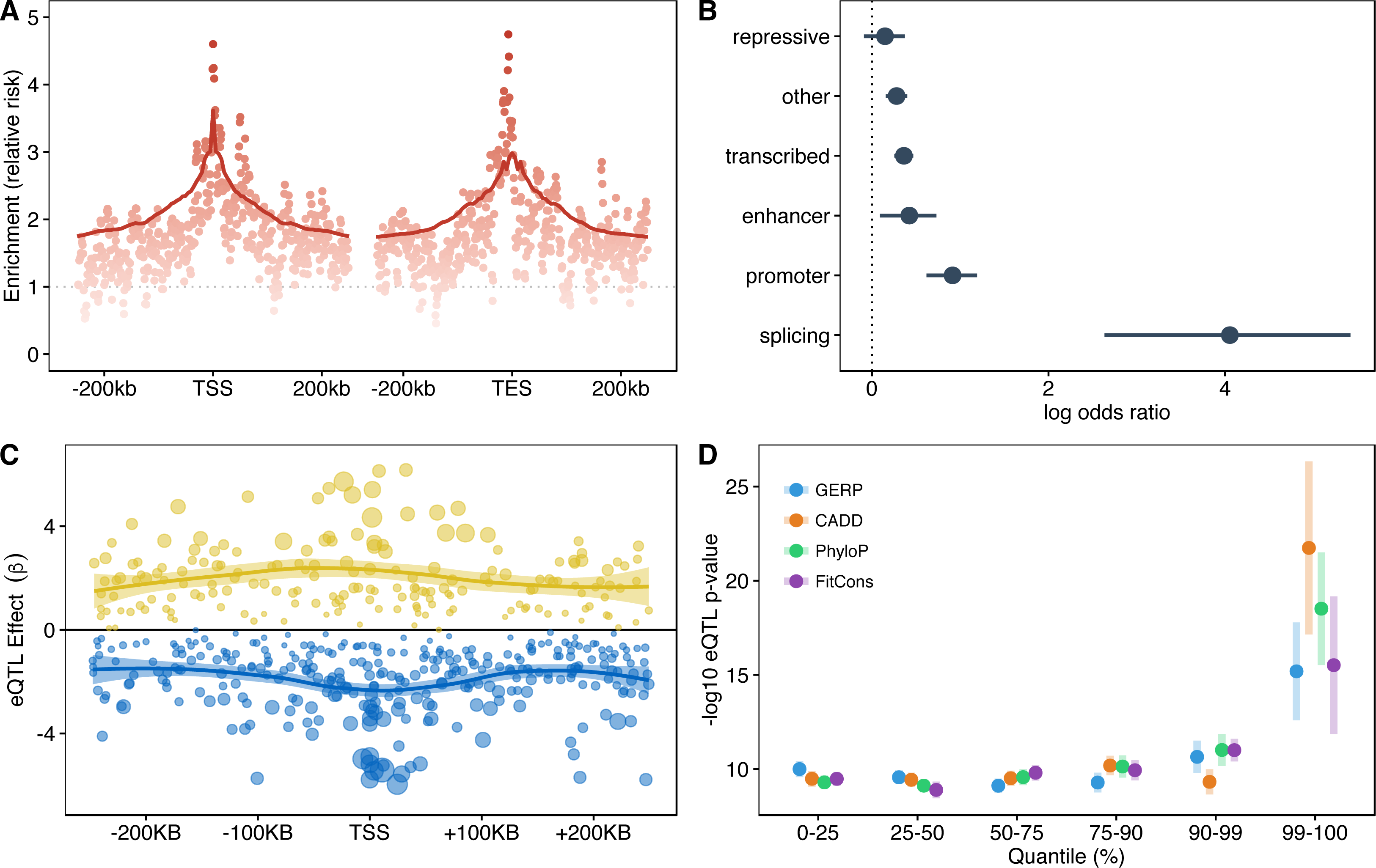
Properties of rare, shared variants near outlier genes. **(a)** Relative enrichment in the number of rare variants transmitted between outlier parents and children versus non-outlier parents and children. Relative enrichments were calculated in overlapping windows of 5KB for the 250KB regions adjacent to the TSS and TES of outlier genes. Enrichment is measured as the relative risk of finding rare shared variants in outlier versus non-outlier lineages in each window. **(b)** Shared rare variants in outlier lineages are enriched for functional regions of chromatin in peripheral blood and splice donor/acceptor regions. **(c)** The position of shared rare variants is plotted relative to the TSS against the regression coefficient derived from the rare eQTL analysis. The color represents under-expression (blue) and over-expression (yellow) rare eQTLs, and the size indicates relative significance. **(d)** Metrics of conservation, evolutionary constraint, fitness, and deleteriousness can identify the most significant rare eQTLs.

Based on these observations, we developed a strict set of rules to distinguish putatively causal rare variants by prioritizing variants that were close to the TSS or likely involved in splicing, highly conserved, and replicated their effects in the larger population (see Methods). We identified candidate causal variants for 30 outlier genes (Supplemental Table S13), including five rare splicing variants. One of these splicing variants, rs35933842, is found at the first exon-intron boundary of *P2RX7*, a ligand-gated ion channel responsible for ATP-dependent lysis of macrophages. While rs35933842 is rare in all European populations including Sardinia, where it is most frequent with a MAF = 0.009% (Supplementary Table S12), it has been previously been shown to disrupt proper splicing of *P2RX7*, leading to an elongated transcript that is subsequently degraded by nonsense-mediated decay and results in mono-allelic expression (Skarratt et al., 2005). As expected, all carriers of rs35933842 (n = 12) in the Sardinia cohort under-expressed *P2RX7* and all reads showed the same allele. While the other splicing variants have not been characterized, we saw similar trends for all five splicing variants suggesting that all of these putative splicing variants are effectively null alleles (Figure 5).

**Figure 5.**
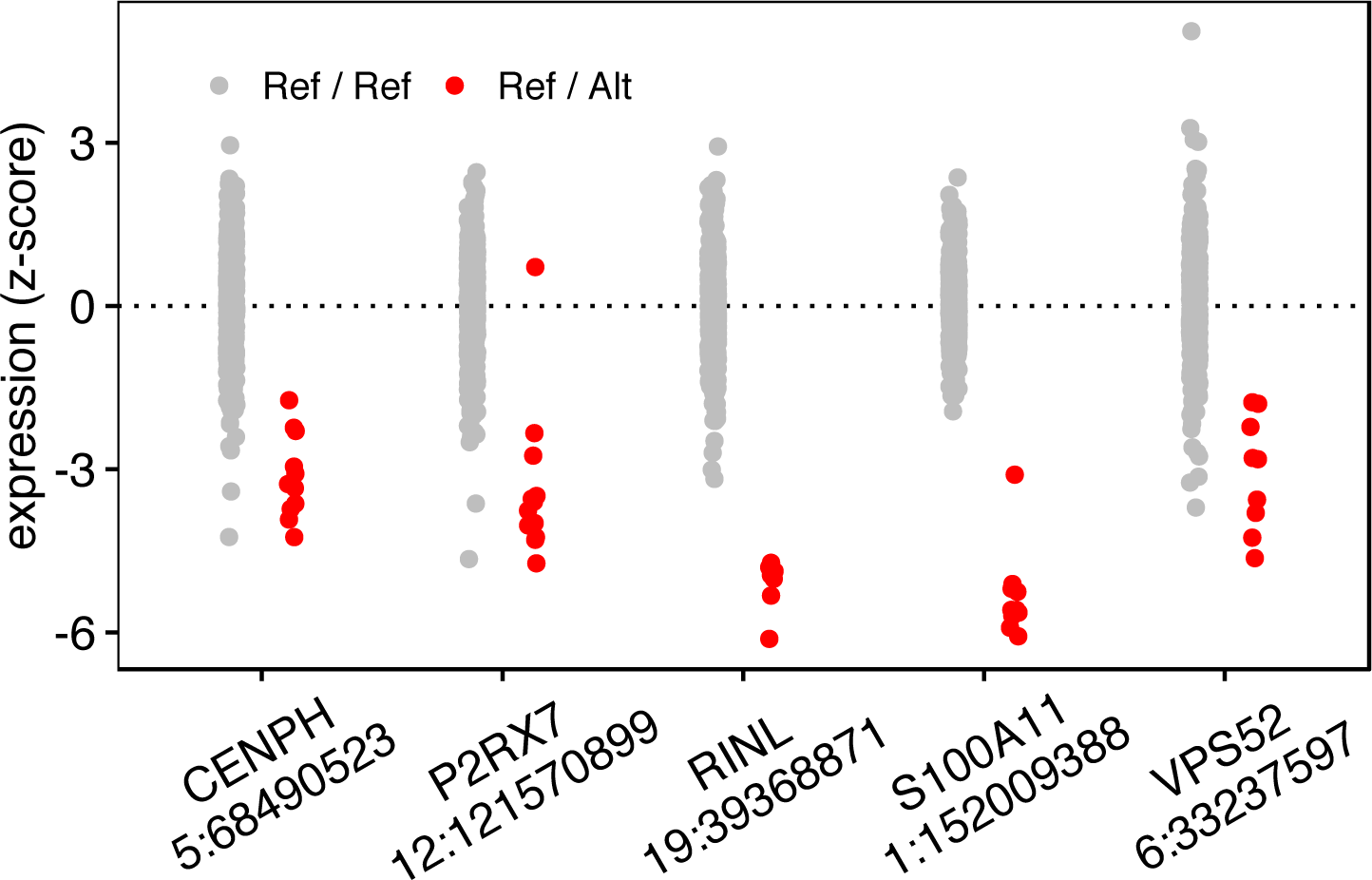
Gene expression patterns in carriers of rare splicing variants. We identified five splicing variants in under-expression outliers - for each variant, the expression level of the affected gene is shown in red for heterozygous carriers in the Sardinia cohort and gray for individuals homozygous for the reference allele.

Because the SardiNIA cohort has been extensively phenotyped, we were able to test for the association of rare variants with measured traits. Of the 30 putatively causal variants, 11 were associated with the expression of genes near significant GWAS loci. Of these, five genes (*SPECC1*, *GLB1*, *CADM1*, *BRI3BP*, and *ANXA5*) were associated with traits measured in the Sardinia cohort. However, we found no significant association between the five candidate variants for these genes and their matched GWAS traits (Supplementary Table S14). Furthermore, we found no significant relationship between expression levels of these genes and their matched GWAS trait (Supplementary Table S14), suggesting that either the gene is not involved in the trait or that dosage is not a critical factor. We next searched for outlier genes that have established roles in the manifestation of rare clinical traits. We were able to identify three outlier genes associated with clinical traits in our database: *VPS13D* is known to repress interleukin-6 (IL6) production; *TSSC1* suppresses osteolysis; and mutations in *POMGNT1* disrupt dystroglycan and can interfere with skeletal muscle function. For each gene, we tested the genotype of the candidate rare variant with levels of the appropriate trait and then for the overall association between gene expression and the trait (see Methods). We were unable to find any significant evidence for association, consistent with recent observations in British-Pakistani cohorts for association testing of rare protein-coding variants in trait-associated genes (Johnston et al., 2015; Narasimhan et al., 2016). While we were unable to identify any direct association between rare variants and clinical traits, we did observe a modest enrichment of outliers in potential disease genes and a marked enrichment of outlier genes in loss-of-function intolerant genes relative to common eQTLs (Supplemental Figure S9).

## DISCUSSION

In our study, we focused on identifying the effect of population and individual-specific non-coding variants in Sardinia. By including secondary QTLs identified through conditional analyses, we identified thousands of variants associated to both expression and splicing. Several hundred eQTLs were completely novel in Sardinia including an eQTL for *ARHGDIB* that was also associated to changes in neutrophil percentage. By comparing Sardinian and European eQTLs, we identified signatures of the Sardinia founder effect and selection on eQTL variants. We also observed an enrichment of eQTLs with large allele frequency differences for multiple sclerosis and malaria, both important factors in the island’s epidemiologic history. We next took advantage of whole genome sequencing, transcriptomes, and familial relationships to identify and describe rare variants with large effects on gene expression. We focused on patterns of outlier gene expression to implicate the functional role of rare regulatory variants (Li et al., 2014; Montgomery et al., 2011; Zeng et al., 2015; Zhao et al., 2016). Such observations have previously been largely limited to unrelated individuals (Zeng et al., 2015; Zhao et al., 2016); but by analyzing segregating patterns of extreme expression in 61 Sardinian trios, we were able to identify hundreds of genes showing large effects and estimated that each individual carries on average 13 genes with an expression outlier (median z-score of 2.97). We implicated rare variants underlying these expression outliers. Furthermore, we observed that outlier expression effects were more prevalent in genes intolerant of loss-of-function variation, consistent with increased potential for important individual consequences. Taken together, these results demonstrate how population history can shape the distribution of common and rare non-coding variants and ultimately influence individual health and disease risk. We anticipate that increasingly large catalogues of functional non-coding variants—found either in isolated populations or families—will yield new opportunities for clinical interpretation, precision health, and the understanding of genome biology.

## ACKNOWLEDGEMENTS

All participants gave informed consent, with protocols approved by institutional review boards for ASL4 in Sardinia and by the University of Michigan. IRB exemption (OHSRP #11916) applied to analyses on coded data at collaborating institutions. Z.Z. is supported by the National Science Foundation (NSF) GRFP (DGE-114747). Z.Z. and J.R.D. also acknowledge support from the National Institute of Health (NIH) (T32HG000044). J.R.D. is supported by the Stanford Graduate Fellowship. K.R.K. is supported by DoD, Air Force Office of Scientific Research, National Defense Science and Engineering Graduate (NDSEQ) Fellowship, 32 CFR 168a. S.B.M. is supported by the National Institutes of Health through R01HG008150, R01MH101814 and U01HG007436. The SardiNIA project is supported in part by the intramural program of the National Institute on Aging through contract HHSN271201100005C to the Consiglio Nazionale delle Ricerche of Italy. S.B.M. would like to thank the SCGPM for computational infrastructure for supporting this project. M.P. would like to thank the CRS4 for sequencing and computational infrastructure for supporting this project.

